# Genos-m: a foundation model for human-associated microbial genomes

**DOI:** 10.64898/2026.05.21.726868

**Authors:** Chao Fang, Fangming Yang, Hao Hou, Huahui Ren, Huanzi Zhong, Huinan Xu, Jiahao Zhang, Jianxin Su, Jielun Cai, Jingnan Yuan, Leo Jingyu Lee, Junhua Li, Kui Wu, Lihui Wang, Liwen Xiong, Long Hou, Meng Ni, Shida Zhu, Shiping Liu, Sirong Liu, Ting Zhu, Xiaofang Chen, Xiaofeng Wang, Zhan Xiao, Xin Jin, Xinting Liu, Xuyang Feng, Yinbin Qiu, Yujing Liu, Yupeng Zhou, Yuxiang Lin, Zhaorong Li, Zhouming Huang, Zhun Shi

## Abstract

Human-associated microbial genomes encode extensive strain-level diversity and niche-specific gene repertoires that are critical to host health. However, these complex sequence features remain difficult to capture using general-purpose DNA foundation models, highlighting the need for dedicated representation learning tailored to the human microbiome. Here, we introduce Genos-m, an open-source foundation model for human-associated microbial genome representation. Genos-m was pretrained on approximately 1.2 trillion nucleotide tokens from a curated microbial genome corpus, including human-associated prokaryotic isolates, high-quality metagenome-assembled genomes (MAGs) and bacteriophages, supplemented with GTDB species-level representative genomes to broaden prokaryotic taxonomic breadth. The model uses a sparsely activated Mixture-of-Experts (MoE) Transformer architecture, with 4.7 billion total parameters, approximately 330 million activated parameters per forward pass and a maximum context length of one million base pairs.

We evaluated frozen Genos-m representations across short-sequence and gene-level tasks, biosynthetic gene cluster (BGC)-based regional sequence tasks, whole-genome strain phenotype prediction, and zero-shot transfer on prokaryote-related RNAfitness assays. Across these benchmarks, Genos-m consistently ranked among the leading comparison models, with the best performance in five of eight gene-fitness regression tasks and in BGC type classification. Using sparse autoencoders, we identified sparse features in Genos-m hidden activations that aligned with annotated ORFs, intergenic regions, and tRNA and rRNA loci.

In downstream applications, Genos-m-derived genome-informed species representations incorporated into a human microbiome self-supervised learning model improved colorectal cancer (CRC)-control classification over conventional species-abundance random forest models. Genos-m also generated stable sample-level embeddings from as few as 10,000 metagenomic reads, retaining gut microbial community structure that distinguished geographic origin and aligned with enterotypes defined from full-depth taxonomic profiles.

Together, these results support Genos-m as a reusable representation model for microbial genomes and metagenomes, with conclusions bounded by the reported datasets, task definitions and evaluation protocols. Genos-m model weights, inference code, and usage documentation are publicly available on GitHub (https://github.com/BGI-HangzhouAI/Genos-m) and Hugging-Face (https://huggingface.co/BGI-HangzhouAI/Genos-m).

## Introduction

Human-associated microbial genomes, which are closely associated with host physiology and health, present a particularly important and technically demanding target for representation learning. They colonize the human gut, oral cavity, skin, respiratory tract and urogenital tract, and are shaped by host genetics, geography and disease states. At the sequence level, they encompass extensive habitat-associated gene repertoires and uncultured diversity captured by MAGs and bacteriophages. Horizontal gene transfer and recombination further redistribute accessory genes and reshape local gene neighbourhoods. These processes continually expand strain-level variations far beyond species labels or representative genomes, leading to substantial inter-individual and habitat-specific heterogeneity.

Current microbiome analyses still rely largely on reference-based taxonomic profiling and gene annotation [7, 8]. These approaches remain essential for interpretable taxonomic and functional compositions, but they are constrained by predefined features and reference coverage [6, 8, 9]. More importantly, they do not provide a reusable sequence-derived representation that connects microbial genes, genomic regions, whole genomes and metagenomic samples across habitats, cohorts and studies.

Genomic foundation models (GFMs) use self-supervised pretraining on vast DNA sequences to learn reusable representations for sequence comparison, search, prediction, and downstream modelling [1, 3, 4]. Recent general-purpose GFMs have demonstrated the utility of broad DNA pretraining, but their corpora are typically optimized for wide taxonomic or cross-domain coverage. In parallel, emerging biological foundation models support the rationale for tailoring pretraining to specific biological sequence spaces. For example, virus-focused models such as LucaVirus [64], which jointly model viral nucleotide and protein sequences across diverse viral sequence space, highlight the value of domain-aligned pretraining for downstream biological applications. This principle is particularly relevant for human-associated microbiomes, where extensive uncultured microbial diversity and genomic variation are central to downstream tasks and underscore the need for targeted exposure to this sequence space during pretraining.

We therefore developed Genos-m, a genomic foundation model tailored to human-associated microbial sequence space. Built on the Genos architecture [2], Genos-m adapts this framework for microbial genome representation learning through targeted corpus construction, sparsely activated expert scaling, and million-base context support. To ensure broad exposure to human-associated microbial diversity, we assembled a stratified pretraining corpus comprising approximately 1.2 trillion nucleotide tokens from high-quality human-associated prokaryotic isolate genomes, MAGs and curated bacteriophage genomes. GTDB species-level representative genomes were further included to broaden prokaryotic taxonomic context.

We evaluate Genos-m as a reusable sequence representation model for microbial genomes and metagenomes. With the pretrained model kept frozen, we assess whether Genos-m-derived embeddings retained task-relevant information across benchmarks covering promoter and gene-level classification, gene-fitness regression, BGC-regional sequence tasks, whole-genome strain phenotype prediction and zero-shot transfer to prokaryote-related RNAfitness assays. We further examined its use in metagenomic applications, including CRC case-control classification and low-input read-based sample representation.

Together, these analyses present Genos-m as a domain-tailored genomic foundation model for human-associated microbes, and examine its utility as a reusable representation backbone across microbial genome and metagenome modelling tasks.

## Results

### A curated corpus supports human-associated microbial genome modelling

The Genos-m pretraining corpus was constructed to represent both broad prokaryotic taxonomic context and human-associated microbial sequence diversity. We first included species-level representative prokaryotic genomes from GTDB R220 to provide broad prokaryotic sequence regularities [5]. We then incorporated public human-associated microbial genomes from UHGG, AWI-Gen2, gcMeta, IMGG, CGR, HROM, PMDB, and related resources [6, 10–14]. To further increase strain-level diversity, we added in-house high-quality MAGs, which contributed approximately one third of the final training tokens after stratified sampling. Phage genomes from UHGV were also included to represent viral components of the human microbiome [15].

After quality control and stratified sampling, the final pretraining corpus contained approximately 1.2 trillion nucleotide tokens and covered 186 phyla, 3,448 families, and 69,056 species. Within this corpus, the retained human-associated prokaryotic subset covered 45 phyla, 585 families, and 12,273 species. Prokaryotic genomes were retained if they were isolate genomes or MAGs with completeness greater than 90% and contamination below 1%. Phage genomes were retained if they were complete or high-quality genomes with greater than 90% completeness according to CheckV. Reverse-complement augmentation was applied with a 50% probability, and context length was progressively extended across 8k, 32k, 128k, and 1M stages.

### Sparse expert scaling supports million-base microbial genome contexts

Genos-m follows the Genos architecture design principle that sparse MoE layers provide modelling benefits beyond computational efficiency [2]. Expert specialization and conditional computation allow the model to draw on a larger parameter space while activating only a subset of parameters for each token. In Genos-m, this design yields 4.7 billion total parameters and approximately 330 million activated parameters per forward pass (Figure 1). Compared with Genos, Genos-m expands the expert pool from 8 to 32 experts and retains Top-2 routing, so the router dynamically selects two experts for each nucleotide token according to sequence content.

**Figure 1:**
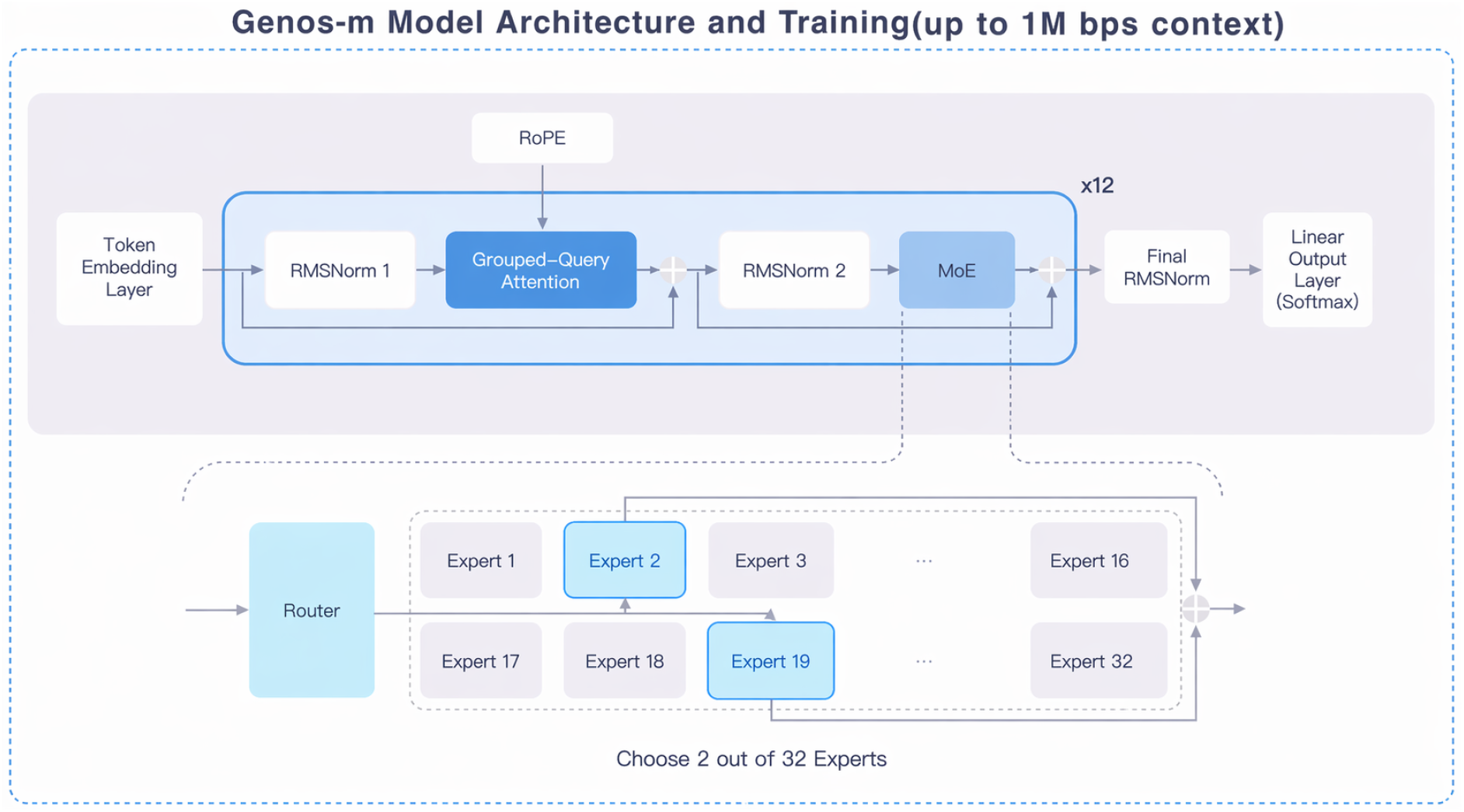
Genos-m model architecture. Genos-m inherits the core MoE architecture and training paradigm of Genos, while expanding the feed-forward expert capacity to 32 experts with Top-2 routing.

Genos-m operates at single-nucleotide resolution, with ambiguous and unknown tokens excluded from the language-modelling loss. As in Genos, the model uses token embeddings, RMSNorm layers, Rotary Position Embedding (RoPE) with a base of 50 million [61], grouped-query attention, and SwiGLU expert feed-forward blocks [62] to support million-token genomic contexts. RoPE injects positional information during attention computation rather than through explicit input position embeddings, and grouped-query attention balances long-context efficiency with representational capacity. During training, expert load balancing, z-loss, gradient clipping, FlashAttention [58], and parallelism across tensor, pipeline, context, data, and expert dimensions were used to improve stability and scalability. During inference, the model retains sparse Top-2 expert routing and FlashAttention-based long-context computation.

This sparse MoE architecture makes Genos-m practical for long-context inference compared with much larger dense models that require substantially heavier GPU deployment. Although the model has 4.7B total parameters, only approximately 330M are activated per forward pass. In inference tests, Genos-m processes approximately 200 kb fragments on 24 GB consumer GPUs, which is sufficient for many bacterial region-level analyses, including metabolic gene clusters, mobile genetic elements and most bacteriophage genomes. On a single 80 GB GPU, it supports 1M bp inference for chromosome-scale or large-contig microbial sequence analysis.

### Genos-m representations support local and regional microbial sequence tasks

We benchmarked Genos-m across short-sequence, gene-level, regional long-context and whole-genome tasks. Unless otherwise specified, evaluations were performed with a frozen pretrained backbone. Final-layer hidden representations were mean-pooled and used to train lightweight task-specific classifiers or regressors under the defined task splits. Classification tasks reported AUROC as the primary metric, with accuracy (ACC) and F1-score as supporting metrics. Regression tasks were evaluated by Pearson correlation.

Across five local sequence and gene-level classification tasks, Genos-m consistently performed in the leading tier among all compared models (Figure 2). It achieved the highest performance for antibiotic-resistance gene (ARG) identification, with an AUROC of 0.9896, ACC of 0.9532, and F1-score of 0.9531. Genos-m also reached AUROC of 0.8460 for promoter prediction, 0.9548 for virulence factor gene (VFG) identification, 0.8534 for essential gene identification, and 0.9932 for six-class bacterial genomic-region classification, covering CDS, pseudogene, rRNA, tRNA, ncRNA and intergenic regions. In these tasks, its performance was comparable to, or ranked second only to, Evo2-40B (Figure 2). These results indicate that frozen Genos-m representations retain task-relevant signals related to short regulatory elements, functional gene categories, and annotated bacterial genomic regions.

**Figure 2:**
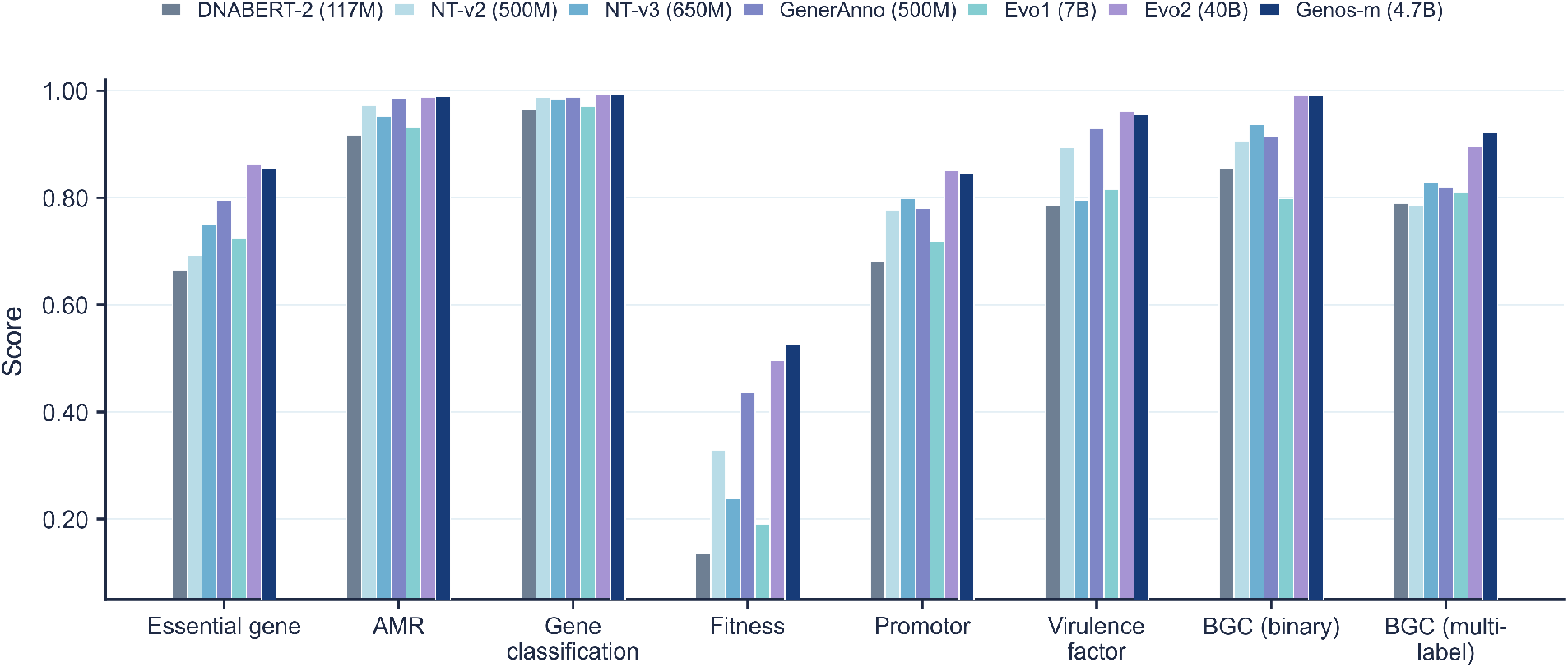
Performance comparison between Genos-m and baseline models across benchmark tasks. Classification tasks are reported using AUROC. Gene-fitness prediction is reported as the mean Pearson correlation across eight regression tasks. Full classification metrics, including AUROC, ACC, and F1-score for each task are provided in Supplementary Table S1. Task-level Pearson correlations for gene-fitness prediction are provided in Supplementary Table S2. Baseline models include DNABERT-2 [43], Nucleotide Transformer [4], Nucleotide Transformer v3 [44], GenerAnno [38], Evo [3], and Evo 2 [1].

The gene-fitness regression benchmark comprised eight condition-specific subtasks, covering both nutrient conditions and chemical stress. Genos-m achieved the best performance among evaluated models in five subtasks: L-arabinose as the sole carbon source, L-histidine nutrient condition, minimal medium with glucose, pyruvate as a carbon source, and perchlorate stress. In the remaining three subtasks, Genos-m ranked second to Evo2-40B. This outperformance on specific nutrient and stress responses suggests that Genos-m’s domain-specialized pretraining provides a crucial advantage over general-purpose massive models in capturing niche-specific ecological adaptations, allowing a highly parameter-efficient model to excel in habitat-specific metabolic and stress-response tasks.

Regional long-context performance was evaluated using two BGC-region tasks. In the BGC versus non-BGC region classification task, complete BGC regions were contrasted with length-matched non-BGC genomic regions. In the BGC class classification task, complete BGC regions were used for multi-label classification across six BGC classes. BGCs provide a representative benchmark for regional microbial genome modelling because recognizing and classifying them requires integrating coordinated genetic features across multiple co-localized genes, despite substantial sequence divergence. In BGC versus non-BGC region classification, Genos-m achieved an AUROC of 0.9907, comparable to Evo2-40B at 0.9911. It also outperformed Nucleotide Transformer v3 and GenerAnno, which reached 0.9365 and 0.9136, respectively. In multilabel BGC type annotation, Genos-m achieved the highest performance among the compared models, with an AUROC of 0.9216. These results suggest that the long-context window and sparse MoE architecture can work together to represent multi-gene, non-local functional modules within candidate genomic regions. They therefore provide direct support for the model’s core design goal of learning microbial genome representations beyond local nucleotide similarity.

### Zero-shot evaluation on prokaryote-related RNAGym RNAfitness assays

We next tested whether a DNA-pretrained microbial genomic model could provide useful zeroshot scores for prokaryote-related RNAfitness assays. We selected 13 RNAGym RNAfitness assays whose identifiers or descriptions indicated a prokaryote origin or relevance [18]. RNA sequences were converted from U to T before scoring. Each mutant sequence was assigned an average next-token log-likelihood score over the full sequence. Within each assay, Spearman correlation was computed between model score and experimental readout. Values of Spearman correlation, AUC and MCC were then averaged across the 13 assays. No task-specific fine-tuning or downstream head training was performed.

Genos-m achieved a mean Spearman correlation of 0.313, with an AUC of 0.653, and an MCC of 0.224 (Table 2). By mean Spearman correlation, it ranked second among the compared models, after Evo2-7B (mean Spearman = 0.390, AUC = 0.691, MCC = 0.284). This result suggests that Genos-m scores can provide informative zero-shot ranking signals for selected prokaryote-related RNA mutant-effect assays, while not implying general RNA foundation-model capability.

**Table 1:** Data sources and composition of the Genos-m pretraining corpus.

| Source | Tokens for training (billion) | Genomes | Dataloader weight (%) | Primary resources |
| --- | --- | --- | --- | --- |
| GTDB R220 | 379 | 67,525 prokaryotic genomes | 31.30 | GTDB R220 |
| Public human-associated prokaryotes | 446 | 102,986 prokaryotic genomes | 36.84 | UHGG, AWI-Gen2, gcMeta, IMGG, CGR, HROM, PMDB |
| Public human-associated phage genomes | 3.8 | 57,514 phage genomes | 0.31 | UHGV genome catalog |
| In-house human gut high-quality MAGs | 382 | 89,320 prokaryotic genomes | 31.55 | In-house data |

**Table 2:** Zero-shot performance on prokaryote-related RNAGym RNAfitness assays. Values are averaged across 13 assays. Best values are shown in bold, and second-best values are underlined. Comparison models include Evo 2 [1], Evo [3], RNAErnie [50], Nucleotide Transformer [4], RNA-FM [52], RiNALMo [51], and GenSLM [45].

| Model | Spearman | AUC | MCC |
| --- | --- | --- | --- |
| Evo2-7B | **0.390** | **0.691** | **0.284** |
| Genos-m | <u>0.313</u> | <u>0.653</u> | <u>0.224</u> |
| Evo1.5 | 0.289 | 0.640 | 0.208 |
| Evo1 | 0.264 | 0.626 | 0.179 |
| RNAErnie | 0.244 | 0.617 | 0.174 |
| Nucleotide Transformer | 0.178 | 0.579 | 0.105 |
| RNA-FM | 0.168 | 0.577 | 0.103 |
| RiNALMo | 0.141 | 0.561 | 0.085 |
| GenSLM | 0.069 | 0.530 | 0.041 |

### Whole-genome embeddings capture bacterial phenotype signals

Whole-genome evaluation used GIDEON-derived strain phenotype labels from BacBench [19]. Genos-m generated whole-genome embeddings directly from nucleotide sequences of each strain, without protein translation or functional annotation. Using genus-disjoint train, validation, and test splits, whole-genome embeddings were evaluated with linear classifiers for Gram status, oxygen tolerance, motility, spore formation, and beta hemolysis. Genos-m ranked among the top-performing models for several labels, and approached protein-derived foundation-model baselines, including ESM-2 and Bacformer (Table 3). These results indicate that nucleotide-level whole-genome embeddings capture global sequence signals associated with several basic bacterial phenotypes. The evaluation does not imply unrestricted phenotype prediction across untested traits or taxa.

**Table 3:** Performance comparison on bacterial phenotype prediction using whole-genome embeddings. Values are AUROC mean *±* SD. Columns denote Gram-positive status, anaerobe status, motility, spore formation, and beta hemolysis. Best values are shown in bold, and second-best values are underlined. Comparison models include DNABERT-2 [43], Nucleotide Transformer [4], ProkBERT [46], ESM-2 [47], ESM-C [48], ProtBERT [49], and Bacformer [40].

| Model | Gram+ | Anaerobe | Motility | Spore | Hemolysis |
| --- | --- | --- | --- | --- | --- |
| Mistral-DNA | 65.58 ± 6.77 | 77.21 ± 2.99 | 66.98 ± 9.54 | 52.59 ± 21.62 | **63.40 ± 15.20** |
| DNABERT-2 | 86.88 ± 5.47 | 88.84 ± 4.07 | 76.09 ± 2.49 | 72.71 ± 16.94 | <u>62.26 ± 16.78</u> |
| NT | 88.12 ± 8.21 | 90.90 ± 1.02 | 81.63 ± 3.84 | 79.25 ± 27.78 | 60.54 ± 16.54 |
| ProkBERT | 97.07 ± 3.41 | 92.50 ± 1.52 | 64.59 ± 5.22 | 83.66 ± 22.84 | 53.06 ± 5.76 |
| ESM-2 | <u>98.48 ± 1.69</u> | 96.14 ± 1.34 | 82.07 ± 4.29 | 89.55 ± 8.27 | 54.80 ± 10.50 |
| ESM-C | **99.73 ± 0.29** | 96.57 ± 2.93 | 74.70 ± 8.69 | 89.91 ± 12.13 | 54.08 ± 6.86 |
| ProtBERT | 97.50 ± 2.39 | 97.92 ± 1.07 | 79.50 ± 6.52 | <u>90.68 ± 7.52</u> | 53.71 ± 12.93 |
| gLM2 | 96.06 ± 4.29 | 91.08 ± 1.84 | 63.98 ± 5.82 | 74.53 ± 30.76 | 50.10 ± 5.57 |
| Bacformer | 97.61 ± 3.16 | **98.92 ± 0.37** | **87.01 ± 9.71** | **90.85 ± 10.38** | 52.05 ± 8.24 |
| Genos-m | 98.41 ± 1.77 | <u>98.14 ± 0.57</u> | <u>83.22 ± 3.52</u> | 90.24 ± 14.58 | 55.50 ± 1.59 |

**Table 4:** Genos-m model parameters.

| Specification | Genos-m 4.7B |
| --- | --- |
| Total parameters | 4.7B |
| Activated parameters | 0.33B |
| Architecture | MoE Transformer |
| Number of experts | 32 |
| Top-k routing | 2 |
| Layers | 12 |
| Attention hidden size | 1024 |
| Attention heads | 16 |
| MoE FFN hidden size | 4096 |
| Vocabulary | 128 with padding token |
| Maximum context length | 1M |

### Sparse autoencoders reveal interpretable genomic structure features

To examine whether Genos-m hidden activations contain interpretable units, we trained sparse autoencoders (SAE) [16, 17] on last-layer activations from approximately 305,000 randomly sampled 32k chunks drawn from the pre-training corpus. Activations were downsampled by one tenth and globally shuffled, yielding a final SAE training set of approximately one billion token-level activations. The SAE used a latent dimension of 4096, Batch-TopK sparsity with *k* = 128, a batch size of 4096, a learning rate 5 × 10^−5^, and one training epoch.

We then searched for sparse features aligned with known genomic structures in the *E. coli* reference genome NC 000913.3. Candidate feature-to-annotation alignment was evaluated in a one-versus-rest setting using a Domain F1-score. High-activation features were identified for ORF, intergenic, tRNA, and rRNA regions (Figure 3). Among ORF-associated features, f2238(+) and f3009(-) showed strand-biased activation patterns, with preferential activation over ORFs annotated on opposite strands (Figure 3). These results provide preliminary evidence that Genos-m internal activations contain separable features corresponding to known genomic structures.

**Figure 3:**
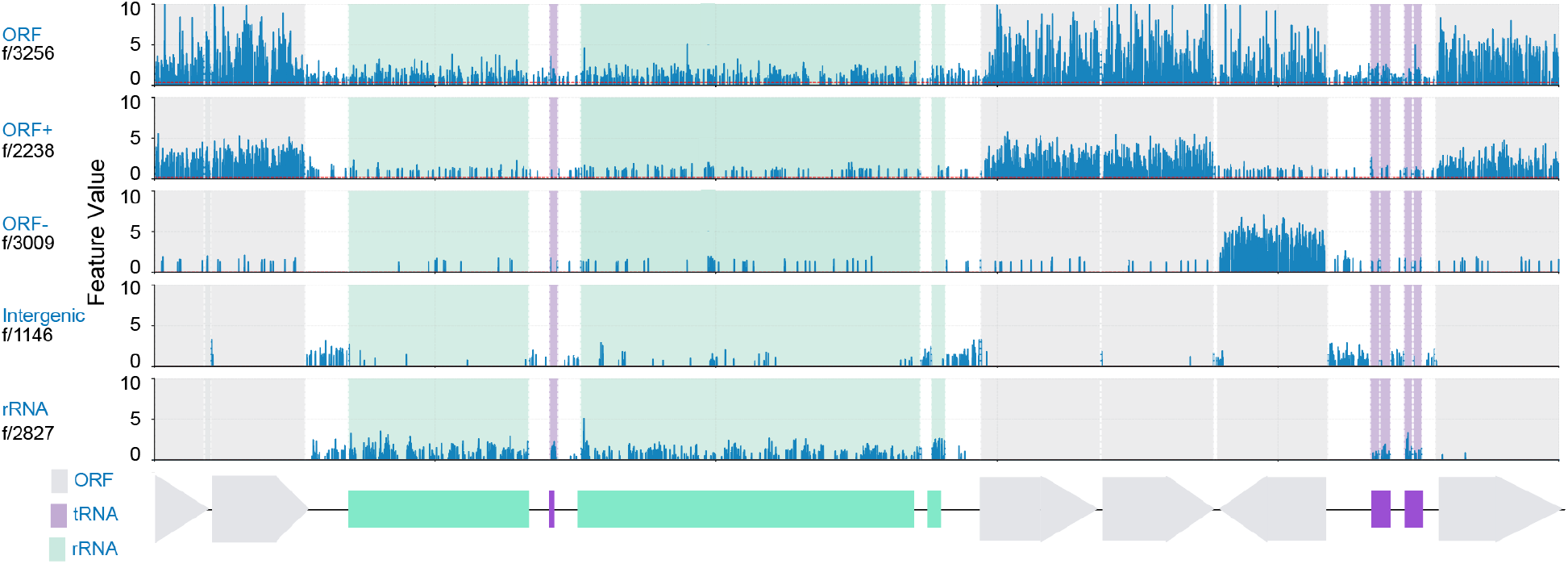
SAE analysis identifies interpretable genomic-structure features in Genos-m activations. Sparse features identified from Genos-m activations align with annotated genomic structures in the *E. coli* reference genome NC 000913.3, including ORF, intergenic, tRNA, and rRNA regions.

### Genome-informed species representations support community-level microbiome self-supervised learning

We applied Genos-m as a genome-representation module for human microbiome community modelling. Genos-m generated genome-level embeddings for representative genomes of gut microbial species detected by MetaPhlAn4 [7]. These embeddings were projected and aligned with species tokens, thereby providing genome-informed species representations for a microbiome self-supervised learning model—a Transformer-based architecture implemented via the *nanoGPT slowrun* framework and pretrained on species-level relative abundance profiles from approximately 400,000 unlabeled human gut shotgun metagenomic samples (Figure 4).

**Figure 4:**
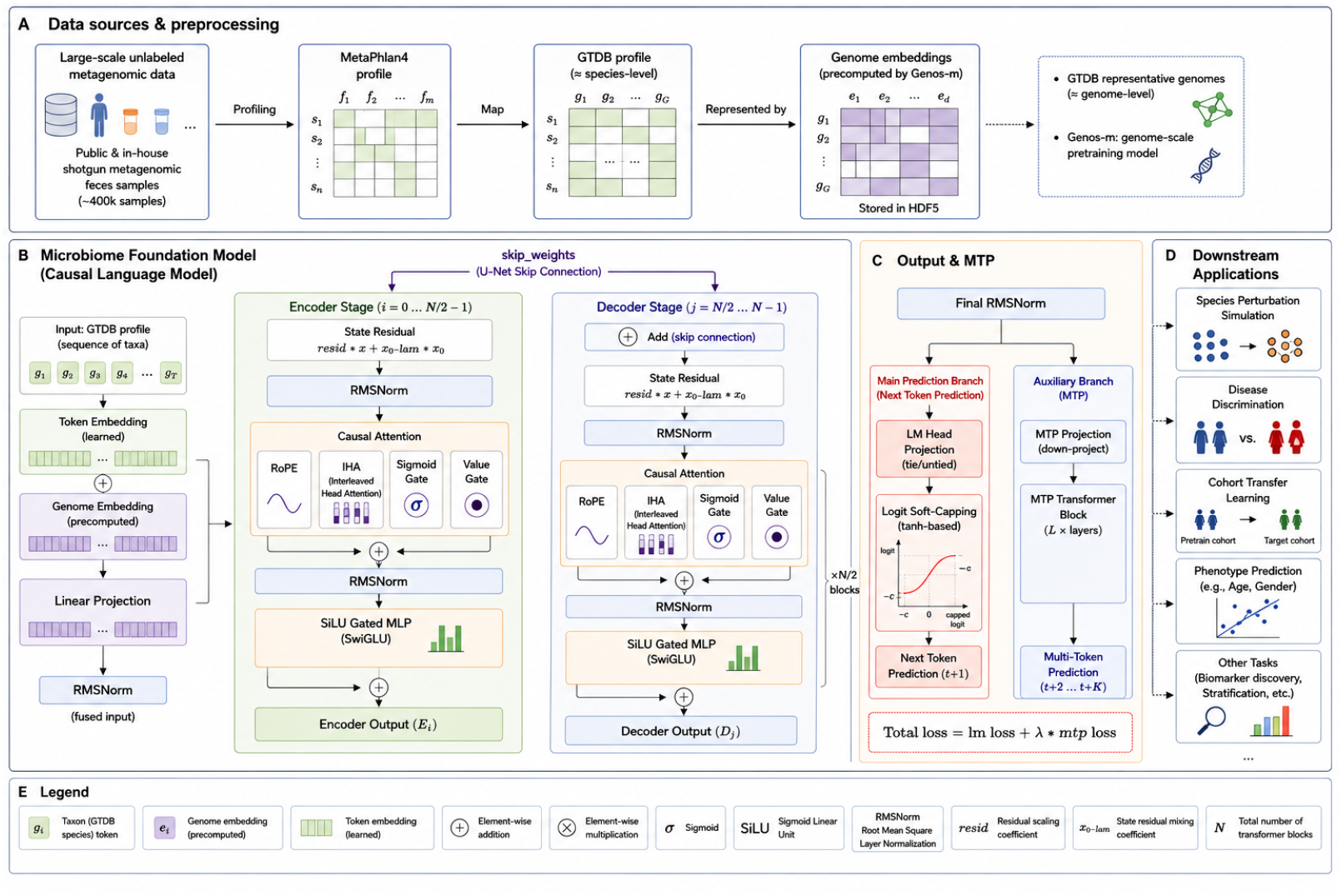
Genome-informed architecture for community-level microbiome self-supervised learning. Representative genome embeddings generated by Genos-m are projected and aligned with species-token embeddings for species-level microbiome representation learning and downstream applications.

Downstream evaluation followed the data and methodology of Piccinno et al. [31], and included 14 independent gut metagenomic cohorts comprising colorectal cancer cases and controls. We compared the genome-informed microbiome self-supervised learning model with the reported species relative-abundance Random Forest (RF) baseline. During classification, the self-supervised backbone was frozen and only a lightweight MLP classification head was trained. In within-cohort 10-fold cross-validation, the genome-informed model achieved a mean AUROC of 0.89, compared with 0.86 for the RF baseline. In pairwise cross-cohort training and testing, it achieved a mean AUROC of 0.77, compared with 0.72 for the RF baseline (Figure 5). These results suggest that Genos-m-derived genome representations can be incorporated into species-level microbiome models while supporting competitive CRC classification performance.

**Figure 5:**
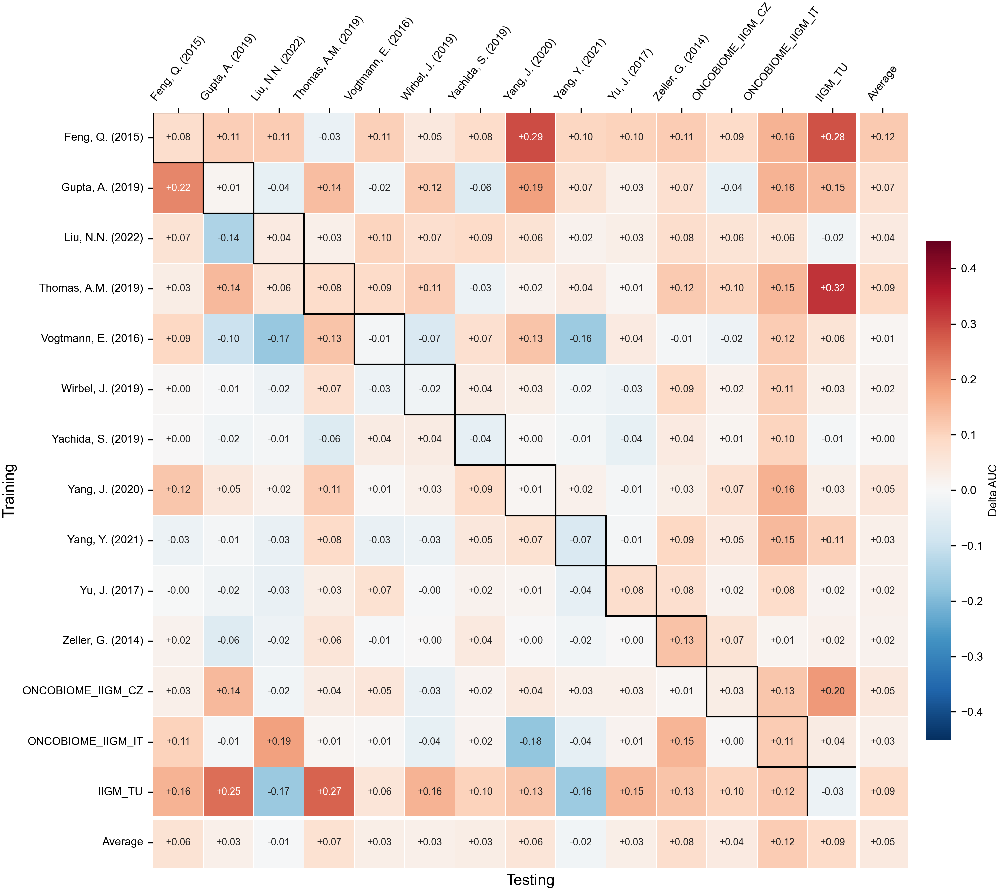
Cross-cohort CRC classification performance between the genome-informed microbiome self-supervised learning model and Random Forest models. Each cell shows the AUROC difference between the genome-informed model and the Random Forest baseline, *AUROC*_model_ − *AUROC*_RF_. Diagonal entries represent within-cohort cross-validation, and off-diagonal entries represent cross-cohort training and testing.

### Low-depth metagenomic embeddings preserve sample structure

We next evaluated whether Genos-m can generate stable sample-level representations directly from low-depth metagenomic reads. Read-level embeddings generated by Genos-m were aggregated into sample-level embeddings by mean pooling. Samples were downsampled to 10K, 100K, 1M, and 10M reads, with five independent replicates generated at each depth.

Across downsampling depths from 10K to 10M reads, embeddings from the same sample clustered tightly in embedding space (Figure 6a). In the two-sample mixture analysis, sample embeddings formed continuous trajectories as mixture proportions changed (Figure 6b). These findings indicate that Genos-m sample-level embeddings are stable across sequencing depth while retaining sensitivity to community-composition perturbation.

**Figure 6:**
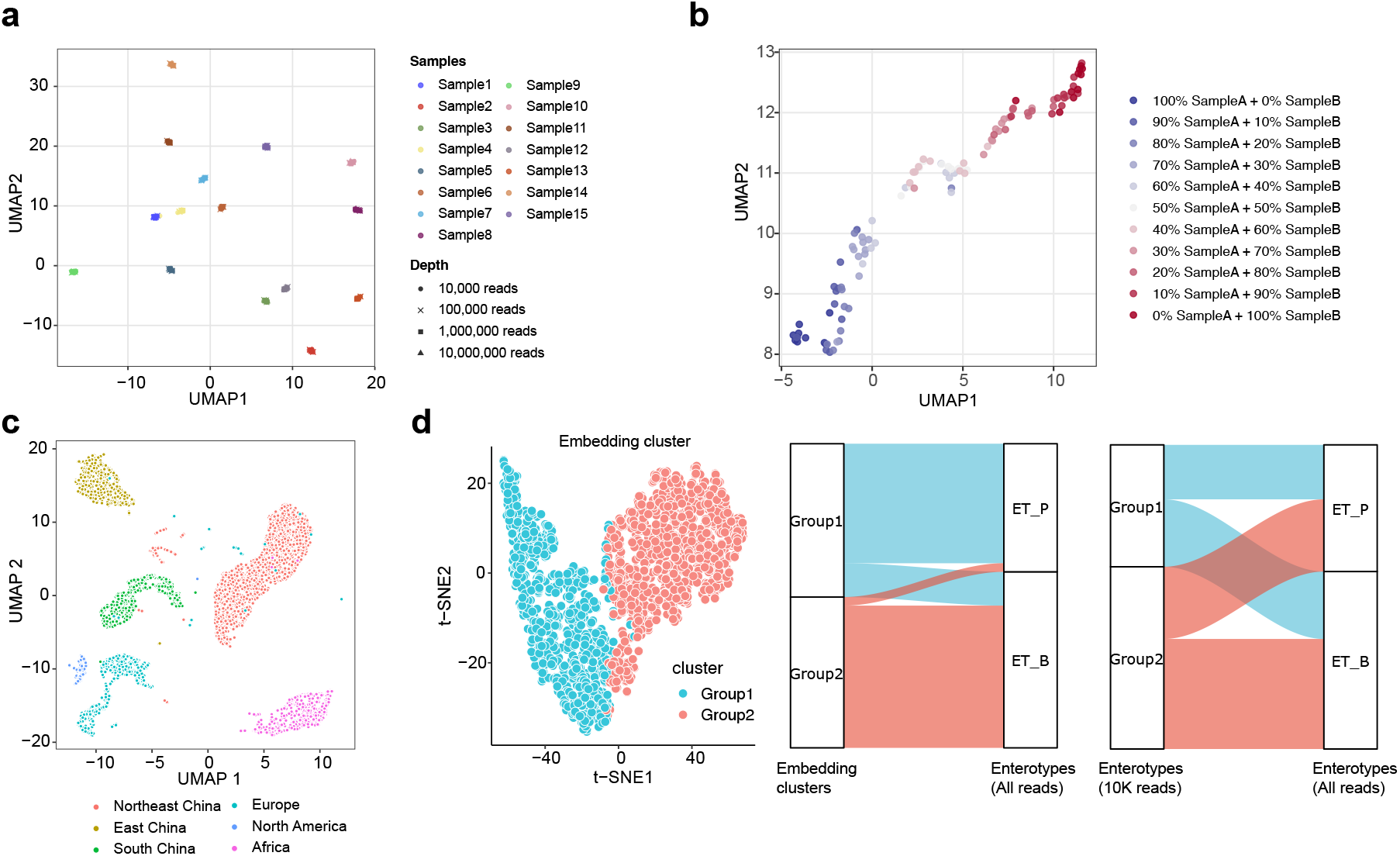
Low-depth Genos-m sample-level representations. Sample-level Genos-m embeddings generated from 10K-read metagenomic inputs show consistency across sequencing depths (a), continuous changes under two-sample mixture gradients (b), separation of host geographic origins (c), and effective recovery of enterotype structure (d).

At the 10K-read setting, low-depth embeddings retained strong host geographic-origin signals. A supervised linear classifier trained on 10K-read embeddings achieved an AUC of 0.998 for multiple geographic-origin classification. Samples from related geographic regions also showed closer organization in embedding space than samples from more distant regions (Figure 6c). Low-depth embeddings additionally recovered enterotype-related structure. Unsupervised clustering of 10K-read Genos-m embeddings formed two clusters that were 86% consistent with reference ET P and ET B enterotypes defined using full-depth genus-level abundance profiles. Under the same 10K-read condition, Kraken2-based genus-abundance clustering showed only 54% consistency with the full-depth reference enterotypes (Figure 6d).

These findings suggest that Genos-m embeddings preserve key sample-level microbiome structure even under ultra-low sequencing depth of 10K reads. Such low-input sample-level representations may support between-sample similarity comparison, host geographic-origin assessment, and quality-control analysis, but do not replace taxonomic profiling.

## Discussion

Genos-m was designed for human-associated microbial genomics, not as a universal DNA model across all domains of life. This distinction matters because prokaryotic genomes are compact, highly variable, and fundamentally different from eukaryotic genomes, with functional variation often distributed across gene neighborhoods, accessory modules, and regional genome organization. Although Genos-m contains substantially fewer activated parameters than very large general-purpose DNA models, such as Evo2-40B [1], it achieved competitive performance across multiple prokaryotic benchmark tasks spanning local sequences, BGC regions, and whole-genome representations. These results support the utility of next-token pretraining on targeted microbial genomes combined with sparse expert activation. The cross-scale consistency further suggests that Genos-m learned more than short-range nucleotide statistics and may capture higher-level regularities in prokaryotic genome organization, thereby enabling reusable representations for microbial genomes and metagenomes.

The whole-genome phenotype benchmarks also highlight a central readout challenge for long-context genome representations. In this study, simple mean pooling of Genos-m final-layer hidden states already captured genome-level signals for several bacterial phenotypes. However, million-base context support does not guarantee that all relevant information is retained in a single fixed-dimensional embedding. Key biologically informative variants, accessory genes, and genomic regions may be diluted by global aggregation. Future improvements will therefore likely require both longer-scale genome modelling and better readout strategies. Longer contexts remain important because typical bacterial genomes can exceed 1 Mbp; chunked whole-genome embeddings can therefore lose cross-chunk dependencies. At the same time, better aggregation is needed to preserve biologically informative sequences.

The RNAfitness task provides a useful boundary case. Genos-m showed informative zero-shot scoring on selected prokaryote-related RNAfitness assays after U-to-T conversion, indicating a degree of transferability from DNA pretraining to RNAfitness scoring in this benchmark subset. This result does not make Genos-m a general RNA foundation model. A more appropriate interpretation is that, in prokaryotes, DNA sequence, transcript sequence, and coding constraints are tightly coupled. A DNA-pretrained microbial model may therefore partially recover RNA-related signals when these constraints are reflected in genomic sequence.

The microbiome applications illustrate a broader role for Genos-m in extending microbiome analysis beyond abundance-based profiles by providing sequence-derived representations at both species and sample levels. Existing microbiome foundation models often treat species as discrete tokens and primarily learn abundance and co-occurrence structure [32, 33], making it difficult to represent genome- or strain-level functional differences. By incorporating genome-derived continuous embeddings, Genos-m can be integrated into community-level microbiome self-supervised learning (SSL) to provide genome-informed species representations. In the CRC use case, the Genos-m-integrated microbiome-SSL framework outperformed the reported species-abundance Random Forest baseline in both within-cohort cross-validation and cross-cohort transfer settings. This result supports the feasibility and downstream utility of genome-informed species representations for microbiome disease modelling. Low-depth metagenomic embeddings suggest that Genos-m can construct sample-level representations directly from raw reads, without first requiring taxonomic assignment or abundance estimation. Whereas conventional taxonomic profiling requires sufficient read depth to quantify taxon abundances reliably, 10K-read Genos-m embeddings retained key sample-level structure across sequencing depths, and recovered geographic-origin signals and enterotype-like structure. This may reflect that low-depth reads provide sparse but distributed samples of community content, while related sequence fragments may occupy nearby regions of the learned embedding space. Mean pooling across read-level embeddings can therefore approximate a sample-level sequence signature. Such low-input representations could enable rapid reference-free sample comparison, cohort stratification, and population-scale prescreening, particularly where full-depth taxonomic profiling is impractical.

Overall, Genos-m establishes a sequence-derived representation framework for microbial genomes and metagenomes. In this study, its utility is demonstrated primarily through frozen representations, including embeddings for genomic sequences, whole genomes and metagenomic samples. These representations provide a complementary layer to annotation-based microbial genomics and establish a basis for exploring additional model-derived signals. Future work should test Genos-m across broader microbial ecosystems, including host-associated, marine, soil and other environmental niches, while improving aggregation methods for local, whole-genome and metagenomic representations. Beyond final embeddings, hidden states, attention patterns, intermediate-layer representations and generative probabilities may encode complementary signals at multiple biological scales. These signals could support feature-level probing, state-based interpretation, retrieval over informative genomic regions, latent perturbation and carefully validated sequence-generation workflows.

More broadly, Genos-m should be viewed as an adaptable genomic representation infrastructure rather than only as a static feature extractor. Continued pretraining could update the model as new microbial genomes, metagenomes and ecological contexts become available. Supervised fine-tuning (SFT) could provide a complementary route for steering model behaviour toward defined biological tasks, including phenotype prediction, functional annotation and strain-level analysis. Together, these transfer-learning modes may move microbial genome modelling beyond fixed representations toward controlled updating and task-guided adaptation. Realizing this potential will require careful task design, independent validation, and explicit safeguards against overinterpreting retrieved, perturbed, or generated sequences. Within these boundaries, Genos-m provides a foundation for adaptable computational workflows in microbial function, ecology, evolution and design.

## Methods

### Pretraining data curation and corpus design

The Genos-m pretraining corpus was designed to capture both broad prokaryotic genomic diversity and high-resolution human-associated microbial variation. The corpus included prokaryotic genomes and phage genomes from public and in-house resources, with an emphasis on genome quality, human-associated microbial niches, and population-specific gut microbial diversity.

Prokaryotic and phage genomes were collected from three major sources. First, species-level representative prokaryotic genomes from GTDB R220 were included to provide broad taxonomic coverage across global microbial diversity [5]. Second, human-associated microbial genomes were incorporated from public and curated resources, including commensal prokaryotic isolates, high-quality MAGs, pathogenic bacterial genomes, and human gut phage genomes. Public resources included UHGG, AWI-Gen2, gcMeta, IMGG, CGR, and HROM [6, 10–14], providing coverage across global populations and multiple human body sites, including gut, oral, skin, and reproductive tract microbiomes. Curated human-associated pathogenic bacterial genomes were collected from PMDB, based on NCBI Pathogen Detection resources [63]. Human gut phage genomes from UHGV [15] were incorporated to extend the corpus beyond cellular prokaryotes and improve representation of phage genomic sequence space. Third, to increase representation of population-level gut microbial diversity, we included high-quality gut MAGs from in-house data.

To reduce the impact of assembly incompleteness and contamination on pretraining, all genomes underwent stringent quality control. Prokaryotic isolate genomes and MAGs were retained only when estimated completeness was greater than 90% and contamination was below 1%, as assessed by CheckM [53]. This contamination threshold is more stringent than the commonly used 5% cutoff for high-quality MAGs. Phage genomes were retained if classified by CheckV [54] as complete circular genomes or as high-quality genomes with greater than 90% completeness.

After quality control, dynamic stratified sampling was applied separately for each context length to balance taxonomic breadth, human-associated niche coverage, and representation across data sources, ecological niches, and species. Independent sampling schemes were used for the 8K, 32K, 128K, and 1M context-window corpora.

The final pretraining corpus covered 186 phyla, 3,448 families, and 69,056 species, comprising 67,525 GTDB species-level representatives (31.3%), publicly available human-associated genomes (37.17%; 102,986 prokaryotes and 57,514 phage genomes), and 89,320 in-house human gut-derived MAGs (31.53%) (Supplementary Figure 1). By source niche, human-associated prokaryotic genomes were distributed across gut (88.9%), oral (4.3%), skin (1.5%), female vaginal tract (0.5%), and PMDB-derived pathogenic genomes (4.7%). By geographic origin, 60.5% of genomes were from Asia, with the remaining genomes distributed across Europe (14.5%), North America (6.2%), Africa (6.0%), Oceania (1.0%), South America (0.2%), and unknown regions (11.5%). The retained human-associated genomes covered 45 phyla, 585 families, and 12,273 species, and more than 75.6% of retained genomes had estimated completeness above 95%.

**Supplementary Figure 1.**
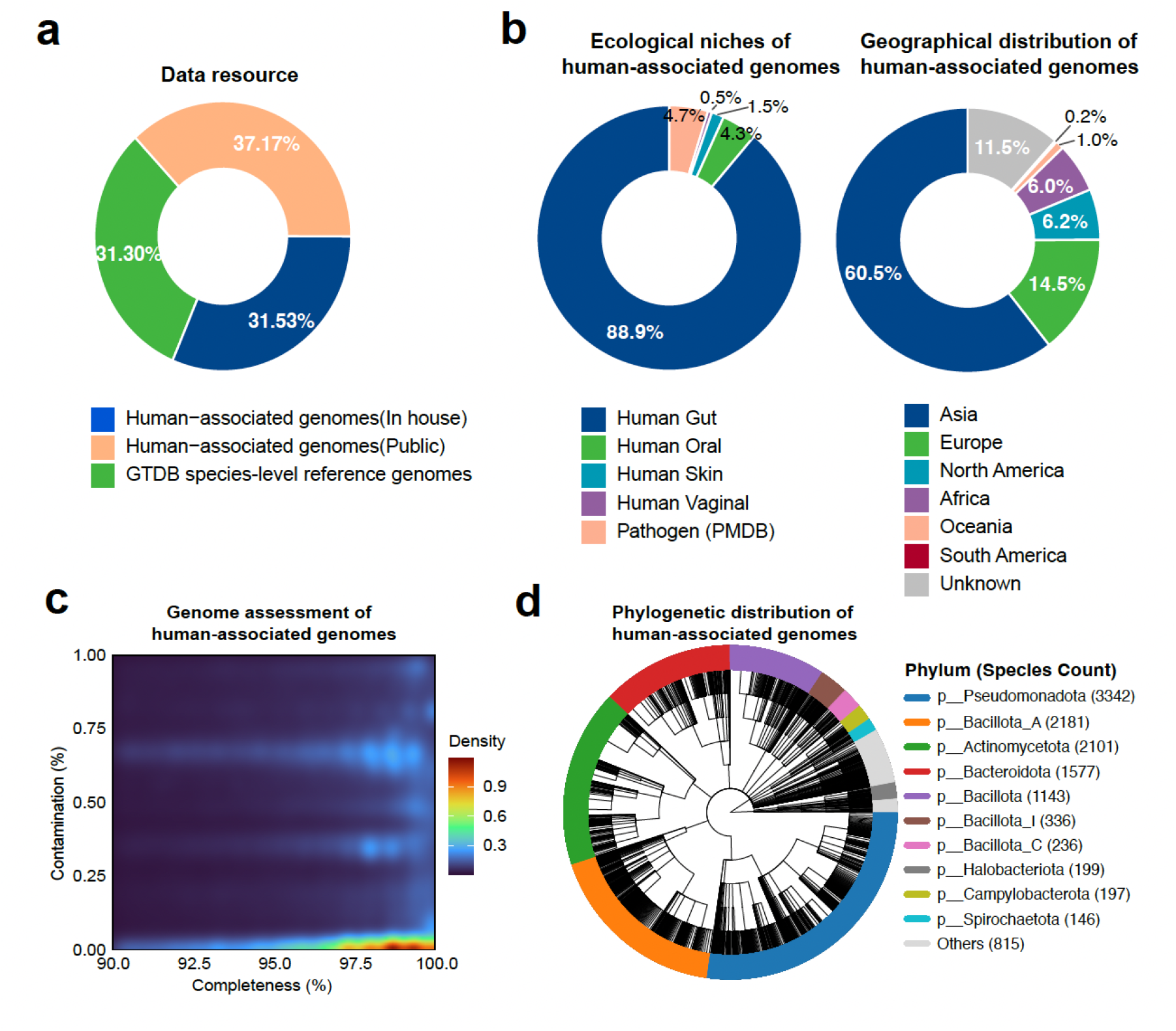
Composition and quality summary of the Genos-m pre-training corpus. Distribution of human-associated prokaryotic genomes across data sources (a), ecological niches and geographic origins (b), genome quality metrics (c), and phylogenetic composition (d).

For each genome, contigs were concatenated using the special separator token # to preserve contig boundaries. Runs of more than three consecutive ambiguous bases (N) were replaced with the special token @ to reduce the influence of long ambiguous regions. Reverse-complement augmentation was applied with a probability of 0.5 during training. The final training corpus contained approximately 1.2 trillion nucleotide tokens.

### Model architecture and training

Bacterial genomes span extensive phylogenetic and functional diversity across taxa, habitats and functional modules. We therefore used a sparsely activated MoE architecture to make long-context modelling tractable while allowing capacity to be allocated conditionally across heterogeneous microbial sequences. Single-nucleotide resolution was retained to reduce the predefined segmentation assumptions imposed by k-mer tokenization and to allow motifs, coding structure, regulatory signals and higher-order sequence units to be learned directly from the data. Long-context modelling was introduced as a biological modelling requirement rather than a scale-driven design choice, because many bacterial traits depend on coordinated gene neighbourhoods, mobile elements and broader genomic organization. Extending the context window from 8K to 32K, 128K and 1M bp allowed sequence units ranging from operons and local gene neighbourhoods to larger genomic islands and most phage genomes to be processed within a single forward pass, when sequence length permitted. Staged context expansion further provided a curriculum-like progression, moving from local sequence regularities, such as codon usage and promoter motifs, toward longer-range gene co-occurrence, neighbourhood structure and broader genome organization.

Genos-m uses a sparsely activated MoE Transformer. The model has 4.7 billion total parameters, 12 layers, attention hidden size of 1024, 16 attention heads, MoE feed-forward hidden size 4096, 32 experts, Top-2 expert routing, and approximately 330 million activated parameters per forward pass. The tokenizer operates at single-nucleotide resolution with a vocabulary size of 128. The model was pretrained with a next-token prediction objective over nucleotide sequences. Ambiguous bases (N) and unknown tokens were excluded from the language-modelling loss. Rotary positional embeddings (RoPE) with a base of 50 million were used to support long contexts [61].

The architecture and training recipe followed the Genos design where applicable [2]. In brief, discrete nucleotide tokens were embedded into continuous representations, stabilized with RMSNorm layers, and processed with RoPE-enabled grouped-query attention. The attention module used 16 attention heads with 8 key-value groups, and each MoE block routed each token to the top two selected expert subnetworks. Expert feed-forward modules used SwiGLU activations [62]. Training used Megatron-LM [57] with tensor, pipeline, context, data, and expert parallelism. Following the Genos training protocol, optimization used AdamW with a cosine learning-rate schedule, a 5% warm-up phase, peak learning rate of 1 × 10^−4^, gradient clipping at 1.0, weight decay of 0.1, and a global batch size of 1,024 with micro-batch size 1. Expert load-balancing auxiliary loss and router z-loss were both set to 1 × 10^−3^. Long-context capability was obtained through progressive context expansion, scheduled learning-rate decay, and RoPE-based context-window scaling. Mixed precision used BF16 for most computations while retaining FP32 precision for numerically sensitive operations, including attention softmax, gradient accumulation and all-reduce communications, and MoE routing.

### Inference hardware and performance

Inference hardware requirements were evaluated using Genos-m 4.7B under single-GPU settings. On one 80 GB GPU, the model weights occupied approximately 9 GB of GPU memory and supported inference with a 1M bp context window. On 24 GB consumer-grade GPUs, Genos-m supported inference for sequence fragments of approximately 200 kb. This context range is sufficient for many local microbial genomic analyses, including metabolic gene clusters, mobile genetic elements, gene-neighbourhood queries, and most bacteriophage genomes.

**Supplementary Figure 2.**
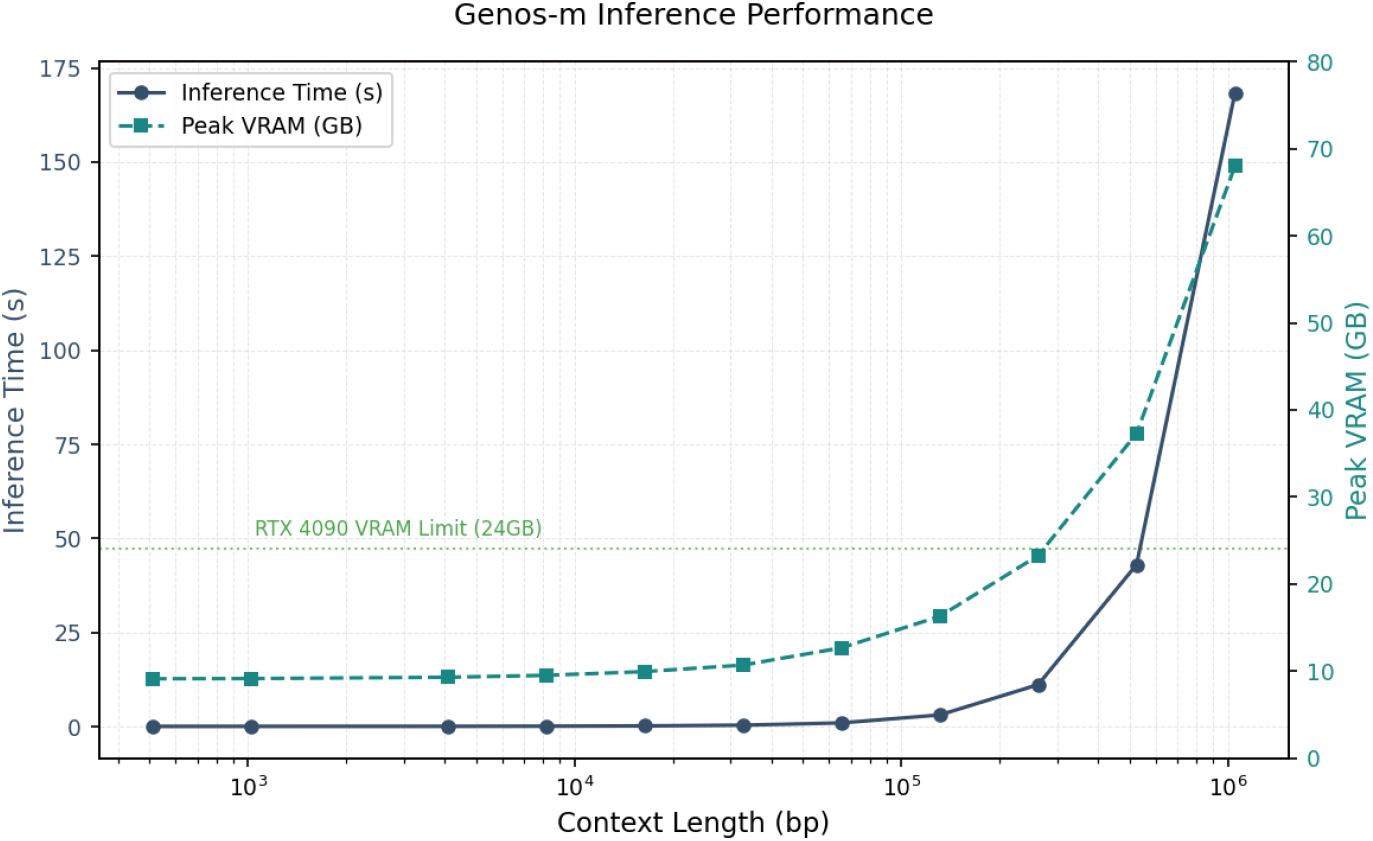
Inference hardware and context-length performance of Genos-m. Inference performance was evaluated under single-GPU settings. Genos-m supports 1M bp context inference on one 80 GB GPU and approximately 200 kb sequence-fragment inference on 24 GB consumer-grade GPUs.

### Unified supervised evaluation

For supervised benchmark tasks, pretrained model backbones were frozen. Input sequences were passed through each model to extract hidden representations. Unless otherwise stated, final-layer token representations were mean-pooled to obtain sequence-level embeddings. Lightweight MLP heads were then trained for classification or regression. Classification tasks used AUROC as the primary metric and also reported ACC and F1-score. Regression tasks used Pearson correlation as the primary metric.

### Local sequence tasks

Essential gene identification used the BacBench essential-gene DNA task [19]. The train, validation, and test partitions followed BacBench, with genomes from the same genus restricted to one split. Promoter identification followed the iPro-MP prokaryotic promoter benchmark [39] and used 81-bp windows. Six-class bacterial genomic-region classification reproduced the GenerAnno bacterial gene classification task [38], with six labels: CDS, pseudogene, tRNA, rRNA, ncRNA, and intergenic regions. Antibiotic-resistance gene (ARG) detection reproduced the GenerAnno drug-resistance prediction setting [38], using CARD resistance genes as positives and length-matched non-AMR RefSeq sequences as negatives. Virulence-factor gene (VFG) detection used VFDB core genes [41] as positives and curated non-virulence essential genes as negatives.

### Gene-fitness regression

The gene-fitness benchmark followed the GenerAnno gene-fitness prediction setting [38]. Eight subtasks were selected from Fitness Browser [59] and covered nutrient-source changes and chemical stress: minimal media glucose, L-arabinose carbon source, pyruvate carbon source, D-alanine nitrogen source, ammonium chloride nitrogen source, L-histidine nutrient condition, cisplatin stress, and perchlorate stress. MSE was used as the training loss, and Pearson correlation was reported.

### RNAfitness zero-shot evaluation

The RNAGym RNAfitness subset was filtered to 13 prokaryote-related assays based on assay identifiers or descriptions [18]. The selected subset contained 54,384 mutant sequences. RNA sequences were converted from U to T before scoring. Whole-sequence next-token average log-likelihood was used as the model score for each mutant. Spearman correlation between model scores and experimental DMS or fitness readouts was calculated within each assay. Following the direction-independent RNAGym convention, absolute Spearman was averaged across assays. No supervised training or downstream prediction head was used.

### BGC regional tasks

BGC binary classification used experimentally validated MIBiG v4.0 BGCs as positive examples [42]. BGC entries from fungal or taxonomically unknown sources, as well as abnormal-length sequences, were removed. Negative examples were sampled from BGC-free contigs after antiSMASH screening [55], with length and CDS-count distributions matched to the positive examples. The final dataset contained 2,107 positives and 2,107 negatives.

BGC multilabel type classification used MIBiG v4.0 BGC regional sequences and merged BGC types into PKS, NRPS, ribosomal/RiPP, saccharide, terpene, and other. The dataset was split into train, validation, and test sets at 8:1:1.

### Whole-genome phenotype evaluation

Whole-genome phenotype prediction used GIDEON strain phenotype data from BacBench [19]. The dataset included 1,175 genomes and five binary phenotype labels: Gram positive, oxygen tolerance, motility, spore formation, and beta hemolysis. Genomes were split by genus into train, validation, and test sets at 7:1:2. Whole-genome embeddings were generated by Genos-m and evaluated using independent linear classifiers for each phenotype. Mean performance and variation were computed across random seeds 1, 2, and 3.

### Sparse autoencoder interpretability

Approximately 305,000 chunks of length 32k were randomly sampled from the model pre-training data. Last-layer Genos-m activations were extracted and downsampled by one tenth. Activations were globally shuffled so that tokens within each 32k SAE training window did not originate from the same raw genomic fragment. The final SAE training set contained approximately one billion token-level activations. The SAE was trained to encode activation vectors into sparse latent features and reconstruct the original activations, using a reconstruction objective with an auxiliary dead-feature penalty.

Batch-TopK sparsity retained the largest *kB* activations across a batch size of *B*, with *k* = 128. The SAE was trained for one epoch using a batch size of 4,096 and a learning rate 5 × 10^−5^. Feature search was performed using annotated ORF, intergenic, tRNA, and rRNA regions in *E. coli* NC 000913.3. Feature-to-annotation alignment was quantified using a one-versus-rest Domain F1-score after sweeping activation thresholds.

### Community-level microbiome self-supervised learning with genome-level representations

For community-level microbiome self-supervised learning, Genos-m generated genome-level embeddings for representative genomes of gut microbial species detected by MetaPhlAn4 [7]. These embeddings were injected by projecting them and summing them with learnable species-token embeddings, yielding genome-informed species representations. The microbiome self-supervised learning model was trained as an in-house Transformer on species-level relative-abundance profiles derived from approximately 400,000 unlabeled human gut shotgun metagenomic samples. The implementation followed slowrun’s two-hour training script, with modifications for microbiome species-abundance sequences. It used abundance-sorted causal language modelling, in which species were sorted by decreasing relative abundance and the model predicted subsequent species from preceding species.

The CRC downstream evaluation used 14 global gut metagenomic cohorts and followed the cohort settings of Piccinno et al. [31]. The self-supervised backbone was frozen, and a lightweight MLP classification head was trained. The evaluation tested disease modelling with within-cohort 10-fold cross-validation and cohort transfer with pairwise cross-cohort external validation.

### Low-depth metagenomic embedding evaluation

We evaluated how many metagenomic reads were sufficient for Genos-m to generate stable sample-level embeddings and whether ultra-low-depth embeddings retained host- and community-level microbiome structure. Raw paired-end reads were preprocessed by reverse-complementing read 2 and concatenating it with read 1 to generate strand-consistent nucleotide inputs. Genos-m generated embeddings for individual reads, which were aggregated into sample-level embeddings by mean pooling.

First, we assessed embedding stability across sequencing depths. Fifteen samples were randomly selected and downsampled to 10K, 100K, 1M, and 10M reads, with five independent replicates generated at each depth. Sample-level embeddings generated from 10K reads were compared with embeddings generated from higher read depths using UMAP visualization and embedding-space consistency analyses. This analysis was used to determine whether 10K reads were sufficient to approximate the sample-level embedding structure obtained from 100K, 1M, and 10M reads.

Second, we evaluated embedding continuity under controlled compositional shifts at the 10K-read setting. For one representative pair of samples, synthetic two-sample mixtures were generated by combining reads from sample A and sample B at 10% increments, from 0% to 100% contribution of sample B. This procedure was repeated ten times. Genos-m sample-level embeddings were then used to assess whether gradual changes in input composition produced continuous trajectories in embedding space.

Third, we tested whether 10K-read embeddings retained geographic-origin signals. This analysis included 4,137 human gut metagenomic samples: 2,033 in-house samples from Northeast China cohorts; 500 publicly available samples from East China cohorts [34]; 500 from a South China cohort [35]; 500 from European cohorts [20, 21, 25, 26, 31]; 500 from African cohorts [36, 37]; and 104 from a North American cohort [22]. Geographic-origin recovery was evaluated at the 10K-read setting using supervised linear classification. Embedding distributions were also examined to assess whether samples from related geographic regions, including cohorts from the same continent, showed closer clustering than samples from more distant regions.

Finally, we evaluated whether 10K-read embeddings retained enterotype structure. In the in-house dataset of 2,033 metagenomic samples, full-depth reads, with an average sequencing depth of 73.6M paired reads, were profiled against the UHGG database [6] using Kraken2 [56] to generate genus-level abundance profiles. Full-depth enterotypes, defined by PAM clustering with Jensen-Shannon distance [60] on genus profiles, were used as the reference, including the Prevotella-dominated enterotype ET P and the Bacteroides-dominated enterotype ET B.

We then generated enterotype assignments from two low-depth representations. For Genos-m, sample-level embeddings generated from 10K reads were clustered using K-means clustering with cosine distance. For the abundance-based baseline, genus profiles generated from 10K-read Kraken2 profiling were clustered using the same PAM-JSD procedure. Low-depth enterotype assignments from both methods were compared against the full-depth enterotype assignments.

## Data availability

Public resources used for training and evaluation are listed in the Methods and References. Genos-m model weights are available on HuggingFace under BGI-HangzhouAI/Genos-m. The Megatron checkpoint is available under BGI-HangzhouAI/Genos-m-Megatron.

## Code availability

Project code, benchmark documentation, and inference examples are available at the Genos-m project repository: https://github.com/BGI-HangzhouAI/Genos-m.

## Acknowledgements

The study was funded and supported by Heilongjiang Provincial Key Research and Development Program (No. SC2024ZX02D0094 to K.W.), National Science and Technology Major Project (No. 2025ZD0551700 to K.W.), and National Natural Science Foundation of China (No. 82570600 to H.R.). The model training process was conducted on the 021 Science Foundation Model and zero2x open platform.

## Author contributions

All authors (C. Fang, F. Yang, H. Hou, H. Ren, H. Zhong, H. Xu, J. Zhang, J. Su, J. Cai, J. Yuan,J. Li, J. Li, K. Wu, L. Wang, L. Xiong, L. Hou, M. Ni, S. Zhu, S. Liu, S. Liu, T. Zhu, X. Chen,X.Wang, X. Liu, X. Feng, Y. Qiu, Y. Liu, Y. Zhou, Y. Lin, Z. Xiao, Z. Li, Z. Huang, Z. Shi) contributed to the design, training, and programming of the Genos-m model. Y. Lin and H. Zhong additionally led the writing and revision of the manuscript. All authors read and approved the final version.

## Supplementary Tables

**Table S1:** MLP classification performance across benchmark tasks. Best values are shown in bold, and second-best values are underlined.

| Task | Metric | DNABERT-2 | NT-v2 | NT-v3 | GenerAnno | Evo1-7B | Evo2-40B | Genos-m |
| --- | --- | --- | --- | --- | --- | --- | --- | --- |
| Essential gene identification | AUC | 0.6658 | 0.6921 | 0.7489 | 0.7953 | 0.7253 | **0.8619** | 0.8534 |
|  | ACC | 0.8704 | 0.8689 | 0.8726 | 0.8863 | 0.8678 | <u>0.8919</u> | **0.8981** |
|  | F1 | 0.5644 | 0.5241 | 0.5452 | 0.6733 | 0.5103 | <u>0.7383</u> | **0.7561** |
| Promoter recognition | AUC | 0.6820 | 0.7773 | 0.7989 | 0.7804 | 0.7185 | **0.8504** | <u>0.8460</u> |
|  | ACC | 0.6937 | 0.7471 | 0.7553 | 0.7502 | 0.7135 | **0.8011** | <u>0.7970</u> |
|  | F1 | 0.6021 | 0.6903 | 0.6990 | 0.6959 | 0.6210 | **0.7679** | <u>0.7562</u> |
| Six-class bacterial gene classification | AUC | 0.9637 | 0.9866 | 0.9848 | 0.9884 | 0.9711 | **0.9937** | <u>0.9932</u> |
|  | ACC | 0.7853 | 0.8887 | 0.8817 | 0.9053 | 0.8026 | **0.9322** | <u>0.9280</u> |
|  | F1 | 0.7821 | 0.8884 | 0.8814 | 0.9052 | 0.7980 | **0.9322** | <u>0.9281</u> |
| Antibiotic-resistance gene detection | AUC | 0.9176 | 0.9727 | 0.9529 | 0.9855 | 0.9313 | <u>0.9884</u> | **0.9896** |
|  | ACC | 0.8274 | 0.9103 | 0.8818 | 0.9440 | 0.8557 | <u>0.9459</u> | **0.9532** |
|  | F1 | 0.8272 | 0.9102 | 0.8817 | 0.9440 | 0.8557 | <u>0.9459</u> | **0.9531** |
| Virulence-factor detection | AUC | 0.7840 | 0.8942 | 0.7939 | 0.9301 | 0.8161 | **0.9617** | <u>0.9548</u> |
|  | ACC | 0.7123 | 0.8086 | 0.6878 | 0.8572 | 0.7385 | **0.8980** | <u>0.8792</u> |
|  | F1 | 0.7116 | 0.8079 | 0.6838 | 0.8568 | 0.7381 | **0.8977** | <u>0.8787</u> |
| BGC binary classification | AUC | 0.8551 | 0.9049 | 0.9365 | 0.9136 | 0.7984 | **0.9911** | <u>0.9907</u> |
|  | ACC | 0.7464 | 0.8081 | 0.8531 | 0.8389 | 0.7180 | <u>0.9076</u> | **0.9289** |
|  | F1 | 0.7358 | 0.8039 | 0.8510 | 0.8378 | 0.6893 | <u>0.9069</u> | **0.9287** |
| BGC multilabel classification | AUC | 0.7897 | 0.7845 | 0.8271 | 0.8207 | 0.8101 | <u>0.8951</u> | **0.9216** |
|  | ACC | 0.2679 | 0.2919 | 0.3206 | 0.3732 | 0.3062 | <u>0.5550</u> | **0.5742** |
|  | F1 | 0.5286 | 0.5242 | 0.5581 | 0.5862 | 0.5257 | <u>0.6512</u> | **0.7384** |

**Table S2:**
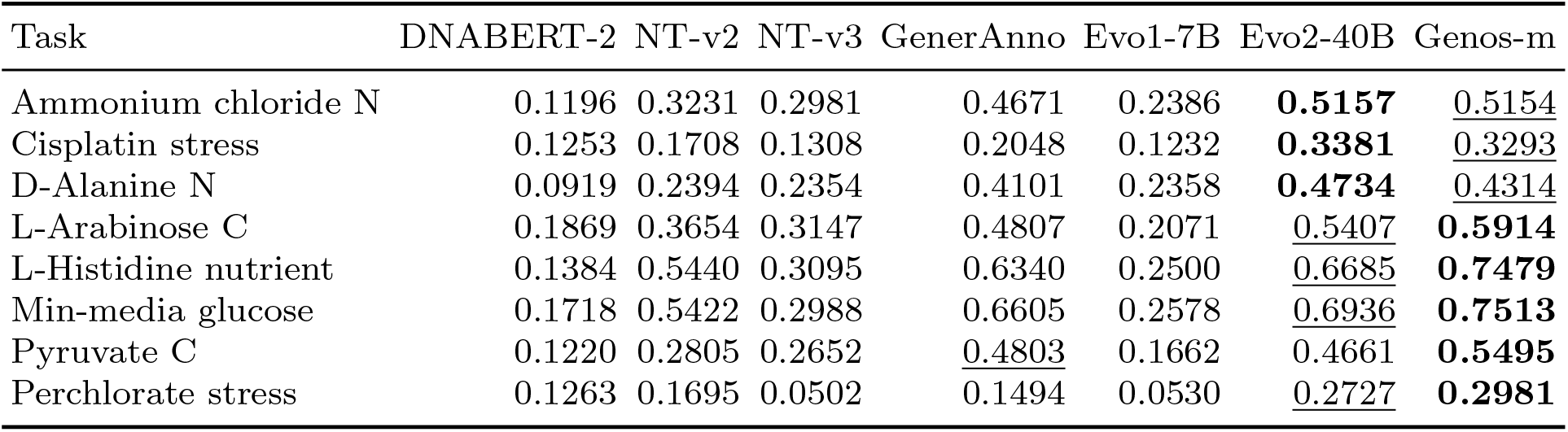
Gene-fitness prediction performance under different environmental conditions. Best values are shown in bold, and second-best values are underlined.

## References

[1] Brixi, G., Durrant, M. G., Ku, J., et al. Genome modelling and design across all domains of life with Evo 2. Nature 652, 1349–1361 (2026).

[2] Lin, A., Xie, B., Ye, C., et al. Genos: a human-centric genomic foundation model. GigaScience 14, giaf132 (2025).

[3] Nguyen, E., Poli, M., Durrant, M. G., et al. Sequence modeling and design from molecular to genome scale with Evo. Science doi:10.1126/science.ado9336 (2024).

[4] Dalla-Torre, H., Gonzalez, L., Mendoza-Revilla, J., et al. Nucleotide Transformer: building and evaluating robust foundation models for human genomics. Nature Methods 22, 287–297 (2025).

[5] Parks, D. H., Chuvochina, M., Rinke, C., et al. GTDB: an ongoing census of bacterial and archaeal diversity through a phylogenetically consistent, rank normalized and complete genome-based taxonomy. Nucleic Acids Research 50, D785–D794 (2022).

[6] Almeida, A., Nayfach, S., Boland, M., et al. A unified catalog of 204,938 reference genomes from the human gut microbiome. Nature Biotechnology 39, 105–114 (2021).

[7] Blanco-Miguez, A., Beghini, F., Cumbo, F., et al. Extending and improving metagenomic taxonomic profiling with uncharacterized species using MetaPhlAn 4. Nature Biotechnology 41, 1633–1644 (2023).

[8] Kieser, S., Brown, J., Zdobnov, E. M., et al. ATLAS: a Snakemake workflow for assembly, annotation, and genomic binning of metagenome sequence data. BMC Bioinformatics 21, 257 (2020).

[9] Bileschi, M. L., Belanger, D., Bryant, D. H., et al. Using deep learning to annotate the protein universe. Nature Biotechnology 40, 932–937 (2022).

[10] Maghini, D. G., Oduaran, O. H., Olubayo, L. A. I., et al. Expanding the human gut microbiome atlas of Africa. Nature 638, 718–728 (2025).

[11] Sun, Y., Chen, Q., Fan, G., et al. gcMeta 2025: a global repository of metagenome-assembled genomes enabling cross-ecosystem microbial discovery and function research. Nucleic Acids Research 54, D724–D733 (2026).

[12] Jin, H., Quan, K., He, Q., et al. A high-quality genome compendium of the human gut microbiome of Inner Mongolians. Nature Microbiology 8, 150–161 (2023).

[13] Zou, Y., Xue, W., Luo, G., et al. 1,520 reference genomes from cultivated human gut bacteria enable functional microbiome analyses. Nature Biotechnology 37, 179–185 (2019).

[14] Cha, J. H., Kim, N., Ma, J., et al. A high-quality genomic catalog of the human oral microbiome broadens its phylogeny and clinical insights. Cell Host & Microbe 33, 1977–1994.e8 (2025).

[15] Camargo, A. P., Baltoumas, F. A., Ndela, E. O., et al. A genomic atlas of the human gut virome elucidates genetic factors shaping host interactions. bioRxiv 2025.11.01.686033 (2025).

[16] Maiwald, A., Jedryszek, P., Draye, F., et al. Decode-gLM: tools to interpret, audit, and steer genomic language models. bioRxiv doi:10.1101/2025.10.31.685860 (2025).

[17] Orlov, A. V., Makus, Y. V., Ashniev, G. A., et al. What do biological foundation models compute? Sparse autoencoders from feature recovery to mechanistic interpretability. bioRxiv doi:10.1101/2026.03.04.709491 (2026).

[18] Arora, R., Angelo, M., Choe, C. A., et al. RNAGym: large-scale benchmarks for RNA fitness and structure prediction. bioRxiv doi:10.1101/2025.06.16.660049 (2025).

[19] Wiatrak, M., Weimann, A., Vinas Torne, R., et al. BacBench: evaluating genomic language models for bacteria. OpenReview, ICLR 2026 withdrawn submission (2025).

[20] Zeller, G., et al. Potential of fecal microbiota for early-stage detection of colorectal cancer. Molecular Systems Biology 10, 766 (2014).

[21] Feng, Q., et al. Gut microbiome development along the colorectal adenoma-carcinoma sequence. Nature Communications 6, 6528 (2015).

[22] Vogtmann, E., et al. Colorectal cancer and the human gut microbiome: reproducibility with whole-genome shotgun sequencing. PLoS ONE 11, e0155362 (2016).

[23] Yu, J., et al. Metagenomic analysis of faecal microbiome as a tool towards targeted non-invasive biomarkers for colorectal cancer. Gut 66, 70–78 (2017).

[24] Gupta, A., et al. Association of Flavonifractor plautii, a flavonoid-degrading bacterium, with the gut microbiome of colorectal cancer patients in India. mSystems 4, e00438–19 (2019).

[25] Thomas, A. M., et al. Metagenomic analysis of colorectal cancer datasets identifies cross-cohort microbial diagnostic signatures and a link with choline degradation. Nature Medicine 25, 667–678 (2019).

[26] Wirbel, J., et al. Meta-analysis of fecal metagenomes reveals global microbial signatures that are specific for colorectal cancer. Nature Medicine 25, 679–689 (2019).

[27] Yachida, S., et al. Metagenomic and metabolomic analyses reveal distinct stage-specific phenotypes of the gut microbiota in colorectal cancer. Nature Medicine 25, 968–976 (2019).

[28] Yang, J., et al. Establishing high-accuracy biomarkers for colorectal cancer by comparing fecal microbiomes in patients with healthy families. Gut Microbes 11, 918–929 (2020).

[29] Yang, Y., et al. Dysbiosis of human gut microbiome in young-onset colorectal cancer. Nature Communications 12, 6757 (2021).

[30] Liu, N.-N., et al. Multi-kingdom microbiota analyses identify bacterial-fungal interactions and biomarkers of colorectal cancer across cohorts. Nature Microbiology 7, 238–250 (2022).

[31] Piccinno, G., et al. Pooled analysis of 3,741 stool metagenomes from 18 cohorts for cross-stage and strain-level reproducible microbial biomarkers of colorectal cancer. Nature Medicine 31, 2416–2429 (2025).

[32] Zhang, H., Zhang, Y., Kang, Z., et al. MGM as a large-scale pretrained foundation model for microbiome analyses in diverse contexts. Advanced Science 13, e13333 (2026).

[33] Pope, Q., Varma, R., Tataru, C., David, M. and Fern, X. Learning a deep language model for microbiomes: the power of large scale unlabeled microbiome data. bioRxiv doi:10.1101/2023.07.17.549267 (2023).

[34] Wu, C., Yang, F., Zhong, H., et al. Obesity-enriched gut microbe degrades myo-inositol and promotes lipid absorption. Cell Host & Microbe 32, 1301–1314.e9 (2024).

[35] Jie, Z., Liang, S., Ding, Q., et al. A transomic cohort as a reference point for promoting a healthy human gut microbiome. Medicine in Microecology 8, 100039 (2021).

[36] Lokmer, A., Cian, A., Froment, A., et al. Use of shotgun metagenomics for the identification of protozoa in the gut microbiota of healthy individuals from worldwide populations with various industrialization levels. PLoS ONE 14, e0211139 (2019).

[37] Tett, A., Huang, K. D., Asnicar, F., et al. The Prevotella copri complex comprises four distinct clades underrepresented in westernized populations. Cell Host & Microbe 26, 666–679.e7 (2019).

[38] Li, Q., Wu, W., Zhu, Y., et al. GENERanno: a genomic foundation model for metagenomic annotation. bioRxiv doi:10.1101/2025.06.04.656517 (2025).

[39] Su, W., Yang, Y., Zhao, Y., et al. iPro-MP: a BERT-based model to predict multiple prokaryotic promoters. Genome Biology 26, 353 (2025).

[40] Wiatrak, M., Vinas Torne, R., Ntemourtsidou, M., et al. A contextualised protein language model reveals the functional syntax of bacterial evolution. bioRxiv doi:10.1101/2025.07.20.665723 (2025).

[41] Zhou, S., Liu, B., Zheng, D., et al. VFDB 2025: an integrated resource for exploring anti-virulence compounds. Nucleic Acids Research 53, D871–D877 (2025).

[42] Zdouc, M. M., Blin, K., Louwen, N. L. L., et al. MIBiG 4.0: advancing biosynthetic gene cluster curation through global collaboration. Nucleic Acids Research 53, D678–D690 (2025).

[43] Zhou, Z., Ji, Y., Li, W., et al. DNABERT-2: efficient foundation model and benchmark for multi-species genome. arXiv preprint arXiv:2306.15006 (2023).

[44] Boshar, S., Evans, B., Tang, Z., et al. A foundational model for joint sequence-function multi-species modeling at scale for long-range genomic prediction. bioRxiv doi:10.64898/2025.12.22.695963 (2025).

[45] Zvyagin, M., Brace, A., Hippe, K., et al. GenSLMs: genome-scale language models reveal SARS-CoV-2 evolutionary dynamics. International Journal of High Performance Computing Applications 37, 683–705 (2023).

[46] Ligeti, B., Szepesi-Nagy, I., Bodnar, B., et al. ProkBERT family: genomic language models for microbiome applications. Frontiers in Microbiology 14, 1331233 (2024).

[47] Lin, Z., Akin, H., Rao, R., et al. Evolutionary-scale prediction of atomic-level protein structure with a language model. Science 379, 1123–1130 (2023).

[48] EvolutionaryScale. ESM Cambrian: revealing the mysteries of proteins with unsupervised learning. EvolutionaryScale (2024).

[49] Elnaggar, A., Heinzinger, M., Dallago, C., et al. ProtTrans: towards cracking the language of life’s code through self-supervised deep learning and high performance computing. IEEE Transactions on Pattern Analysis and Machine Intelligence 44, 7112–7127 (2022).

[50] Wang, N., et al. Multi-purpose RNA language modelling with motif-aware pretraining and type-guided fine-tuning. Nature Machine Intelligence 6, 770–782 (2024).

[51] Penic, R. J., Vlasic, T., Huber, R. G., et al. RiNALMo: general-purpose RNA language models can generalize well on structure prediction tasks. Nature Communications 16, 5671 (2025).

[52] Chen, J., Hu, Z., Sun, S., et al. Interpretable RNA foundation model from unannotated data for highly accurate RNA structure and function predictions. arXiv preprint arXiv:2204.00300 (2022).

[53] Parks, D. H., Imelfort, M., Skennerton, C. T., et al. CheckM: assessing the quality of microbial genomes recovered from isolates, single cells, and metagenomes. Genome Research 25, 1043– 1055 (2015).

[54] Nayfach, S., Camargo, A. P., Schulz, F., et al. CheckV assesses the quality and completeness of metagenome-assembled viral genomes. Nature Biotechnology 39, 578–585 (2021).

[55] Blin, K., Shaw, S., Augustijn, H. E., et al. antiSMASH 7.0: new and improved predictions for detection, regulation, chemical structures and visualisation. Nucleic Acids Research 51, W46–W50 (2023).

[56] Wood, D. E., Lu, J. and Langmead, B. Improved metagenomic analysis with Kraken 2. Genome Biology 20, 257 (2019).

[57] Shoeybi, M., Patwary, M., Puri, R., et al. Megatron-LM: training multi-billion parameter language models using model parallelism. arXiv preprint arXiv:1909.08053 (2019).

[58] Dao, T., Fu, D., Ermon, S., et al. FlashAttention: fast and memory-efficient exact attention with IO-awareness. Advances in Neural Information Processing Systems 35, 16344–16359 (2022).

[59] Price, M. N., Wetmore, K. M., Waters, R. J., et al. Mutant phenotypes for thousands of bacterial genes of unknown function. Nature 557, 503–509 (2018).

[60] Arumugam, M., Raes, J., Pelletier, E., et al. Enterotypes of the human gut microbiome. Nature 473, 174–180 (2011).

[61] Su, J., Ahmed, M., Lu, Y., et al. RoFormer: enhanced transformer with rotary position embedding. Neurocomputing 568, 127063 (2024).

[62] Shazeer, N. GLU variants improve transformer. arXiv preprint arXiv:2002.05202 (2020).

[63] National Center for Biotechnology Information. NCBI Pathogen Detection. https://www.ncbi.nlm.nih.gov/pathogens/ (accessed 29 Nov 2025).

[64] Pan, G., Wang, Y., Huang, H., et al. LucaVirus: large-scale virus nucleotide-protein alignment and multimodal modeling. bioRxiv doi:10.1101/2025.06.14.659722 (2025).

